# Non-enzymatically oxidized arachidonic acid regulates mouse spermatogenic cell T-type Ca^2+^ currents

**DOI:** 10.1101/586297

**Authors:** O. Bondarenko, G. Corzo, F.L. Santana, F. del Río-Portilla, A. Darszon, I. López-González

## Abstract

During spermatogenesis, phospholipids and fatty acids (FAs) play an important role both as structural components of spermatogenic cell plasma membranes and as molecular messengers that trigger the differentiation of the male germ cell line. However, spontaneous oxidation of plasma membrane phospholipids and FAs causes a decrease in mammalian fertility. In the present report, we examine the effects of non-enzymatically oxidized arachidonic acid (AA_ox_) on mouse spermatogenic T-type Ca^2+^ currents (I_CaT_) due to their physiological relevance during spermatogenesis. AA_ox_ effects on the biophysical parameters of I_CaT_ were significantly different from those previously reported for AA. AA_ox_ left shifted the I-V curve peak and both activation and steady-state inactivation curves. I_CaT_ deactivation kinetics were slower in presence of AA_ox_ and the time for its recovery from inactivation increased significantly. Therefore, the fraction of inactivated Ca^2+^ channels of spermatogenic cells is increased at voltages where they are usually active. The inhibition of I_CaT_ by AA_ox_ could contribute to the infertility phenotype and to the observed apoptotic state of spermatogenic cells induced by oxidized FAs.

## INTRODUCTION

During mammalian spermatogenesis, the male germ cell line requires an increase in the intracellular Ca^2+^ concentration ([Ca^2+^]_i_), in order to continue with the cellular differentiation that produces mature sperm. The only voltage dependent Ca^2+^ channels thus far documented in spermatogenic cells (SCs) are T-type Ca^2+^ channels recorded as macroscopic currents (Santi *et al*, 1996; Arnoult *et al*, 1996). They are mainly encoded by the low voltage activated Ca^2+^ channel genes Ca_V_3.2 (α_1H_) and Ca_V_3.1 (α_1G_) in a lower extent (Treviño *et al*, 2004). Previous reports have suggested that T-type Ca^2+^ channels could be important to generate synchronized Ca^2+^ oscillations relevant for [Ca^2+^]_i_ elevation and activation of transcriptional factors involved in spermatogenesis (Sánchez-Cárdenas *et al*, 2012).

In addition to [Ca^2+^]_i_ increase, during spermatogenesis, phospholipids and fatty acids (FAs) also play important roles both as structural components of the SCs plasma membrane and as molecular messengers that trigger the first meiotic division within the male germ cell line, like the testicular meiotic-activating sterol (Keber *et al*, 2013). Sertoli cells provide the majority of phospholipids, sterols and FAs during spermatogenesis. The most representative ones, at least in human male gametes are: phosphatidylcholine and phosphatidylethanolamine, cholesterol and desmosterol, palmitic, arachidonic, and docosahexaenoic acids (Alvarez & Storey, 1995; Keber *et al*, 2013). In particular, AA is released from Sertoli cells in response to follicle-stimulating hormone (Jannini *et al*, 1994). These results strongly suggest that FAs originating in Sertoli cells could be part of the signaling mechanisms that modulate spermatogenic cell metabolism, proliferation, differentiation or death in the seminiferous tubule (Paillamanque *et al*, 2016). *De novo* metabolic synthesis of phospholipids and FAs in Sertoli cells is not enough to cover the spermatogenic cell requirements for these products. Thus, besides *de novo* synthesis, carrier mechanisms of different fatty acids by albumin have been proposed; free transport of different FAs salts is another option (Brash, 2001). Both hypothesis could be right because a deficient diet in essential FAs changes the lipid composition of Sertoli and spermatogenic cell plasma membranes in a period of 9-14 days; however, plasma membrane composition of the germinal cell line is completely recovered when essential FAs and lipids are reincorporated to the diet (Marzouki & Coniglio, 1982).

Arachidonic acid (AA) is a 20-carbon omega-6 polyunsaturated fatty acid and one of the main components of cellular membranes. In addition, AA is an important second messenger involved in signaling cascades being the substrate to produce many different metabolites like prostaglandins, leukotrienes etc. (Meves, 2008). This molecule possesses four cis-double bonds, which are the source of its flexibility and keep this pure fatty acid in a liquid state, even at subzero temperatures. AA participates in different cellular processes like cellular proliferation, where it generates a Ca^2+^ signaling cascade and nitric oxide release (Zuccolo *et al*, 2016). In rat pachytene spermatocytes and round spermatids, AA by itself, not its enzymatically-produced metabolites, induces [Ca^2+^]_i_ increases caused by Ca^2+^ release from intracellular stores. This latter effect could be important for male germ line differentiation since this effect is stronger in spermatids than in spermatocytes (Paillamanque *et al*, 2016).

However, the chemical properties of AA make it subject to multiple possible transformations. Their double bonds are quite propense to react with molecular oxygen. This process can happen non-enzymatically, as consequence of oxidative stress, or through the action of any of three different types of oxygenases: cyclooxygenase (COX), lipoxygenase (LOX), and cytochrome P450 (Brash, 2001; Meves, 2008). While the biological effects of AA and its enzymatic oxidation products are widely discussed in literature, those produced by its non-enzymatic oxidation products are mostly unknown (Brash 2001). In the case of mammalian sperm, the spontaneous oxidation of their plasma membrane phospholipids and FAs has been reported to cause a decrease in fertility (Alvarez & Storey, 1995). For instance, loss of human sperm motility due to O_2_ was reported since 1943 by McLeod (rev. in Alvarez & Storey, 1995). In general, oxidized phospholipids within mammalian sperm produce plasma membrane damage (Jones & Mann, 1973, 1976, 1977, Jones *et al*, 1978, 1979), irrespective of the oxidation mechanism either spontaneous or enzymatic (Alvarez & Storey, 1995).

To contend with oxidized phospholipids and FAs, human sperm possess two principal protecting enzymes: superoxide dismutase (Mennella & Jones, 1980; Alvarez *et al*, 1987) and the group of glutathione peroxidase (GPX), glutathione reductase (GSH) and their substrate glutathione (Li, 1975; Alvarez *et al*, 1987). In addition, human sperm also have catalases (Jeulin *et al*, 1989; Zini *et al*, 1993) and phospholipases A2, which contribute by degrading hydroperoxylated phospholipids and FAs (Alvarez & Storey, 1995). As a consequence of their biological relevance, mutations on the above mentioned enzymes or inhibition of GPX by either depletion of reductive substrate GSH by reaction with H_2_O_2_ or direct inhibition of the enzyme with mercaptosuccinate, increases the rate of spontaneous peroxidation in human sperm by 20-fold and increase infertility in humans (Alvarez & Storey, 1989; Zini *et al*, 1993). In addition, chemical agents like nandrolone decanoate, a synthetic anabolic steroid analog of testosterone used in pig cattle; or cyclophosphamide, a chemotherapeutic agent; both increase phospholipid peroxylation, reduce antioxidant activity, produce spermatic abnormality, apoptosis and DNA fragmentation in mammalian testicles (Chabra *et al*, 2014; Mohamed & Mohamed, 2015). On the contrary, natural antioxidants (melatonin or Rosemary oil) protect against testicular damage produced by phospholipid peroxylation (Chabra *et al*, 2014; Türk *et al*, 2016).

Ion channels such as K^+^, Na^+^ and TRP channels are regulated by AA and some of its enzymatically oxidized forms in different cell types (Chen *et al,* 2001; Basora *et al,* 2003; Meves 2008; Wen *et al,* 2012). It has also been shown that AA inhibits N-, L-and T-type Ca^2+^ channels (Xiao *et al,* 1997; Talavera *et al,* 2004a; Roberts-Crowley & Rittenhouse, 2007; Roberts-Crowley & Rittenhouse, 2009). AA can inhibit α_1G_-encoded T-type Ca^2+^ currents directly, by exerting two independent effects: (a) reducing ion channel availability and (b) shifting the voltage dependence of steady-state inactivation to more negative potentials. Additionally, AA can interact with Ca_V_3.1 (α_1G_) channels in both resting and inactivated states and the structural determinants of inactivation which modulate the AA affinity for this Ca^2+^ channel were suggested (Zhang *et al,* 2000; Talavera *et al,* 2004a; Talavera *et al,* 2004b; Roberts-Crowley & Rittenhouse, 2007; Zhang *et al,* 2013; Baeza *et al,* 2015). Consistently, it has been shown that inhibition of AA metabolism does not affect the AA inhibitory effect on T-type Ca^2+^ channels (Talavera *et al,* 2004b). This result indicates that AA directly, but not its enzymatic products, is inhibiting T-type Ca^2+^ currents in cells. On the other hand, the effects of AA metabolites produced by COX, LOX and cytochrome P450 have been explored on different ion channels (Marnett *et al,* 1999; McGiff & Quilley, 1999; Roman, 2002; Basora *et al,* 2003; Meves, 2008; Zuccolo *et al,* 2016). However, a better characterization of oxidized AA effects on T-type Ca^2+^ channels is lacking.

Considering both that AA is an important regulator of mammalian spermatogenesis and that there is incomplete information regarding how itself and its oxidized metabolites regulate ion channels, the goal of the present project was to investigate the effect of non-enzymatically oxidized AA metabolites (AA_ox_) on T-type Ca^2+^ channels in spermatogenic cells.

## RESULTS

### Synthesis of non-enzymatically oxidized arachidonic acid (AA_ox_

Due to the physiological relevance the early stages of FAs oxidation have on spermatogenesis, we obtained and evaluated how oxidized AA products impact spermatogenic cell T-type Ca^2+^ currents.

We generated non-enzymatically oxidized AA products (AA_ox_) by dissolving AA in ethanol and injecting air into the flask, as described in the materials and methods section. After sample preparation, the AA_ox_ products were separated and identified chromatographically using HPLC. Our results show that AA_ox_ products started to appear after 1 h of air exposure and increased their concentration up to 20 h of treatment (Fig. 1). We found that the oxidation procedure lead to the appearance of 3 major peaks with retention times (RT) at 38 (a relatively small peak), 42 and 45 min, which are not present in the fresh AA sample (Fig. 1A, red line). The generation of AA_ox_ products with this strategy was highly reproducible. All separated peaks were re-purified to detect if the drying and dissolving procedures affected the separated compounds. Interestingly, each of the 3 re-purified peaks lead to obtaining p38, in insignificant amounts, and the p42 and p44 fractions in a higher extent (Fig. 1B). These results allowed us to suggest that the two main AA_ox_ peaks (p42 and p44) observed after the oxidation procedure could correspond to two different conformational stages, which could be in equilibrium and continuously produced by non-enzymatic oxidation. This continued oxidation and equilibration process precluded the purification of the individual fractions, and thus the inhibitory effect described was due to the mixture of both AA_ox_ products. In the text that follows we refer to both products as AA_ox_ to simplify naming them.

**Figure 1.**
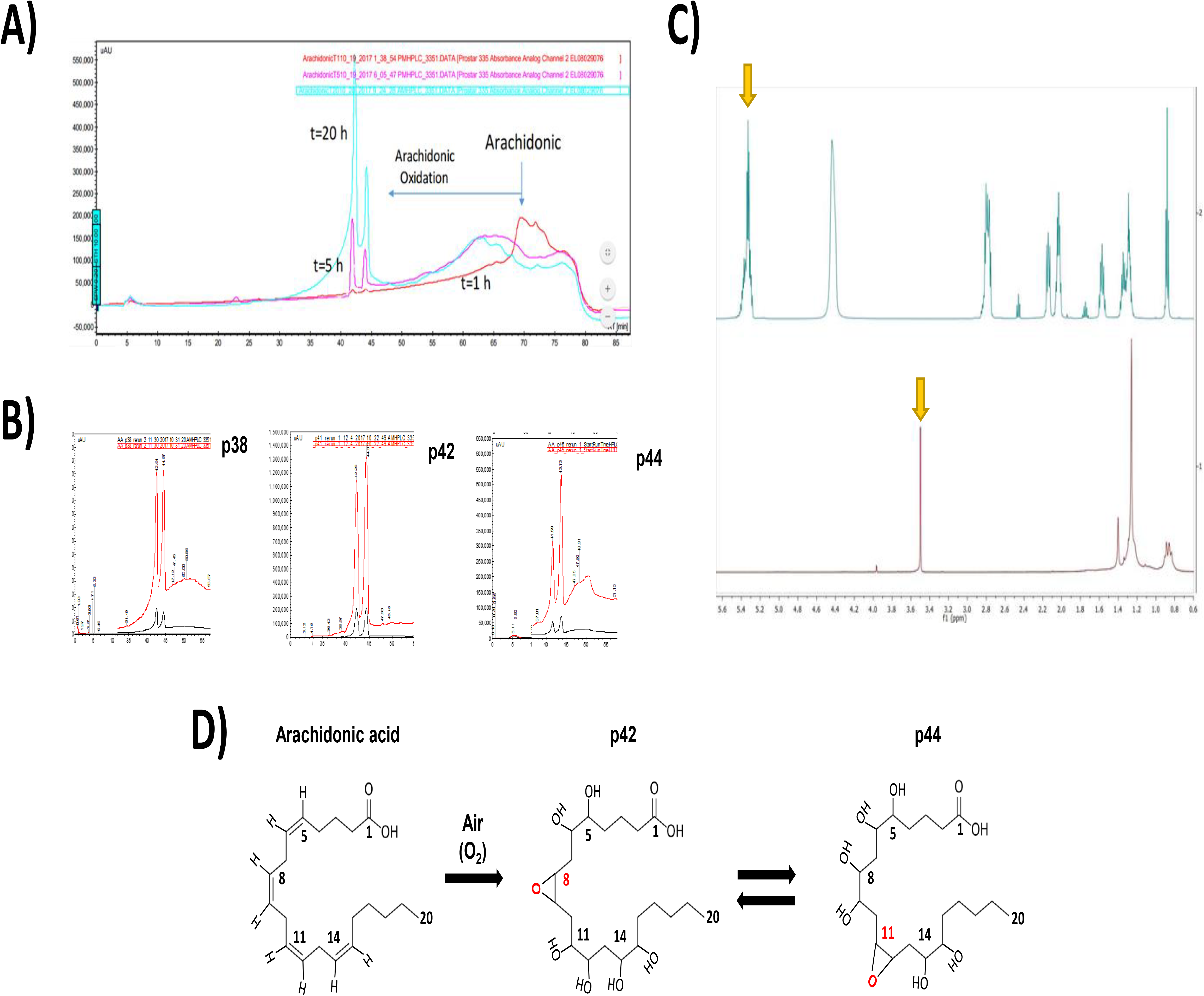
Arachidonic acid (AA) is non-enzymatically oxidized and loses all its double bonds. **A)** Representative RP-HPLC purification of AA_ox_ after non-enzymatic AA oxidation, note that two main fractions are produced after air oxidation during 5 h corresponding to p42 and p45 (pink line). The peak amplitude for each fraction is increased after 20 h of non-enzymatic oxidation of AA (aqua line). A third fraction is also produced in a lower extent, corresponding to p38 fraction. **B)** HPLC re-purification of every independent fraction, namely p38, p42 or p45; reproduced both p42 and p45 main fractions. **C)** Upper panel: representative AA spectrum at 500 MHz, where a signal is observed at 5.3 ppm corresponding to double bond protons (arrows). Lower panel: representative AA_ox_ spectrum at 300 MHz, the signal at 5.3 ppm disappears and one at 3.5 ppm appears which corresponds to CH-OH protons (arrow). These signals and the disappearance of the 5.3 ppm signal are consistent with the oxidation of AA and generation of CH bonded to oxygen. **D)** Structure of arachidonic acid and two of its main non-enzymatically oxidized products in equilibrium (p42 and p44).

^1^H NMR experiments with AA clearly show that the CH from the double bonds at 5.35 ppm (Fig. 1C, upper spectrum). This signal completely disappeared in the ^1^H NMR spectra of the oxidized products and a new signal appears at 3.60 ppm (Fig. 1C, lower spectrum). This result confirms the complete oxidation of AA and the loss of all its double bonds with the formation of base oxygen carbons once the NMR spectrum was acquired.

This analyses plus GC-Ms of p42 and p45 spectra indicated that AA was hyper hydroxylated and completely loss its unsaturated bonds in presence of air, due to AA non-enzymatic oxidation (Fig. 1C). AA_ox_ contained six hydroxyl groups and one epoxide group due to the oxidation of its double bonds. Tautomerism of a hydroxyl and epoxide groups explains the formation of two compounds which interconvert one in each other. The most feasible positions for the epoxide group are double bonds 8 or 11, due to the capacity of interchange this position between them (Fig. 1D). However, the specific structure of AA_ox_ requires additional tests and is currently under study (Fig. EV1).

### AA_ox_ addition shifts the peak of I-V curve to negative potentials

As reported, spermatogenic cells display the classical criss–cross pattern of transitory I_CaT_ (Fig. 2A, control upper traces) (Santi *et al*, 1996; Arnoult *et al*, 1996). Consistently, the I_CaT_ activation threshold was −70 mV and the maximum current amplitude was reached around −37 mV (Fig. 2B, closed circles). Previous reports showed that non-oxidized AA decreases the current amplitude encoded by heterologously expressed Ca_V_3.1 channels (α1G) and does not significantly shift their I-V curve shape (Talavera *et al*, 2004). To examine how non-oxidized AA affects T-type Ca^2+^ currents in mouse spermatogenic cells, peak current amplitudes were measured applying 200 ms voltage pulses from a holding potential (V_h_) of −120 mV up to −40 mV in 5 mV steps. Our results confirmed that AA (5 µM) decreased the I_CaT_ current amplitude of mouse spermatogenic cells around 58% without altering their I-V peak current (Fig. EV2). Surprisingly, addition of a low AA_ox_ concentration (250 nM) inhibited I_CaT_ of spermatogenic cells (Fig. 2A, lower traces; and B opened circles) around 54 ± 6%, and the inhibition extent was stable up to 15 min after AA_ox_ addition (Fig. 2B closed and opened triangles). In addition, the presence of AA_ox_ (250 nM) shifted the peak of the I-V curve around 10 mV to more negative potentials; from −37 ± 1.6 mV, for control, up to −45 ± 0.5 mV, after 10 min of AA_ox_ incubation (Fig. 2B and C).

**Figure 2.**
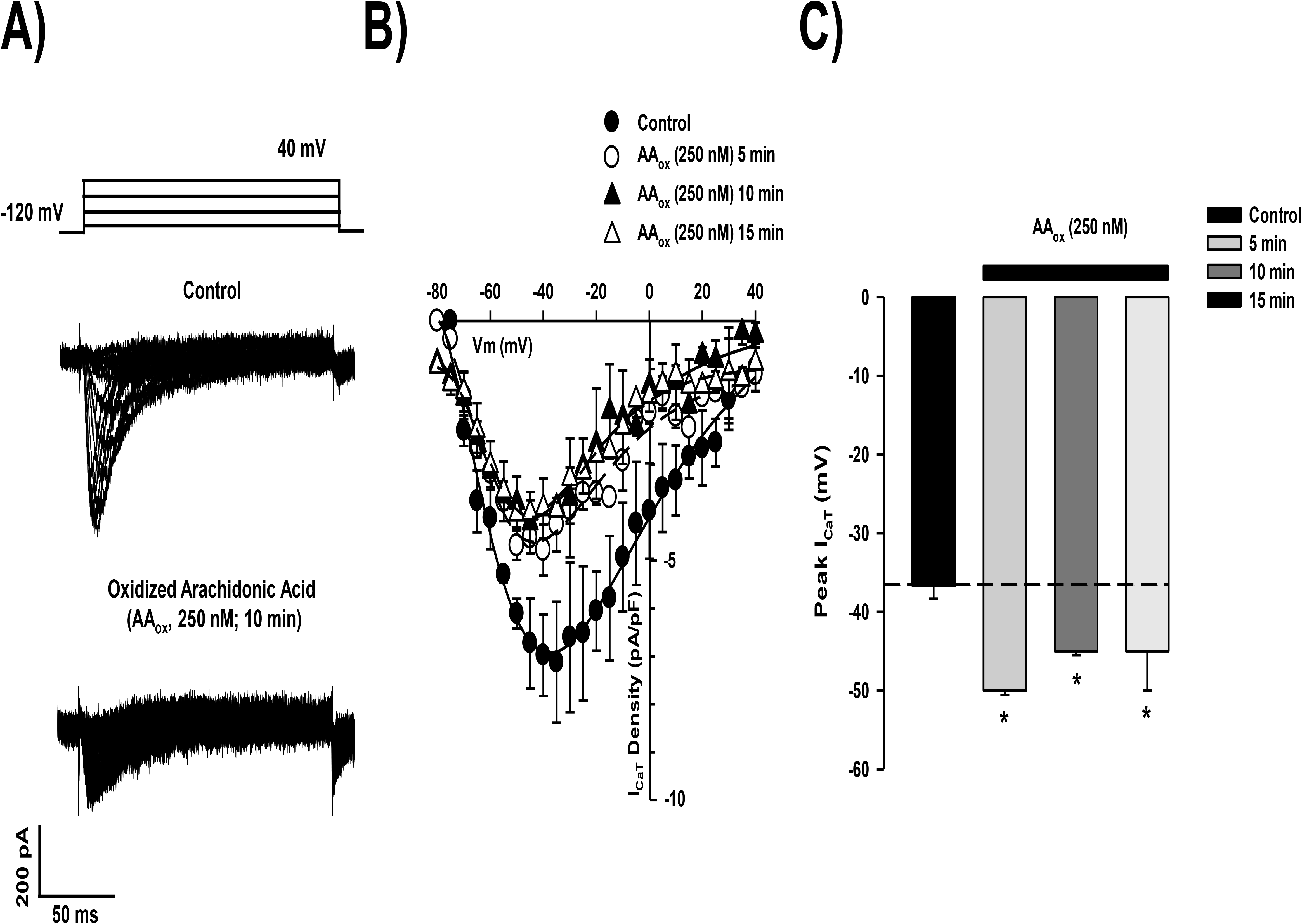
AA_ox_ shifts the I-V curve of spermatogenic cell I_CaT_ toward more negative potentials. **A)** Representative whole-cell T-type Ca^2+^ currents (I_CaT_) of mouse spermatogenic cells in response to 200 ms test pulses of 10 mV steps in the −80 to +40 mV range from a V_h_ of −120 mV, in the absence (control, upper traces) or presence of AA_ox_ (250 nM, lower traces) for 10 min. **B)** Representative current-voltage relationships obtained from the results shown in panel A, but incubating AA_ox_ (250 nM) for different times (5, open circles; 10, closed triangles; or 15 min, open triangles). The I_CaT_ I-V curves peak undergo a left displacement induced by AA_ox_ with respect to the control (closed circles). **C)** Voltage of the I-V curve peak in absence (control: −37 ± 1.6 mV, closed bar) or presence of AA_ox_ (250 nM: −50 ± 0.6, −45 ± 0.5 or −45 ± 5 mV) at different incubation times (5, 10 or 15 min), respectively. In all cases, symbols represent the mean ± S.E.M. (n=4–6).\**p*<0.05

### AA_ox_ is a potent spermatogenic cell T-type Ca^2+^ current inhibitor

In order to confirm our above-mentioned observations, we compared the AA_ox_ inhibitory potency respect to AA-induced inhibition. Currents were elicited with a test pulse at −40 mV from a V_h_ of −120 mV. Representative Ca^2+^ current traces at −40 mV show that 5 and 8 µM AA (Fig 3A, upper panel) and 250 and 1000 nM AA_ox_ (Fig 3A, lower panel) significantly decrease their amplitude. For both AA and AA_ox_, I_CaT_ inhibition was dose dependent; however, the IC_50_ of AA (4.7 ± 0.4 µM) was significantly higher than the one of AA_ox_ (186 ± 12 nM) (Fig. 3B). Hill coefficients were 1.6 ± 0.3 and 1.4 ± 0.1 for AA and AA_ox_, respectively.

**Figure 3.**
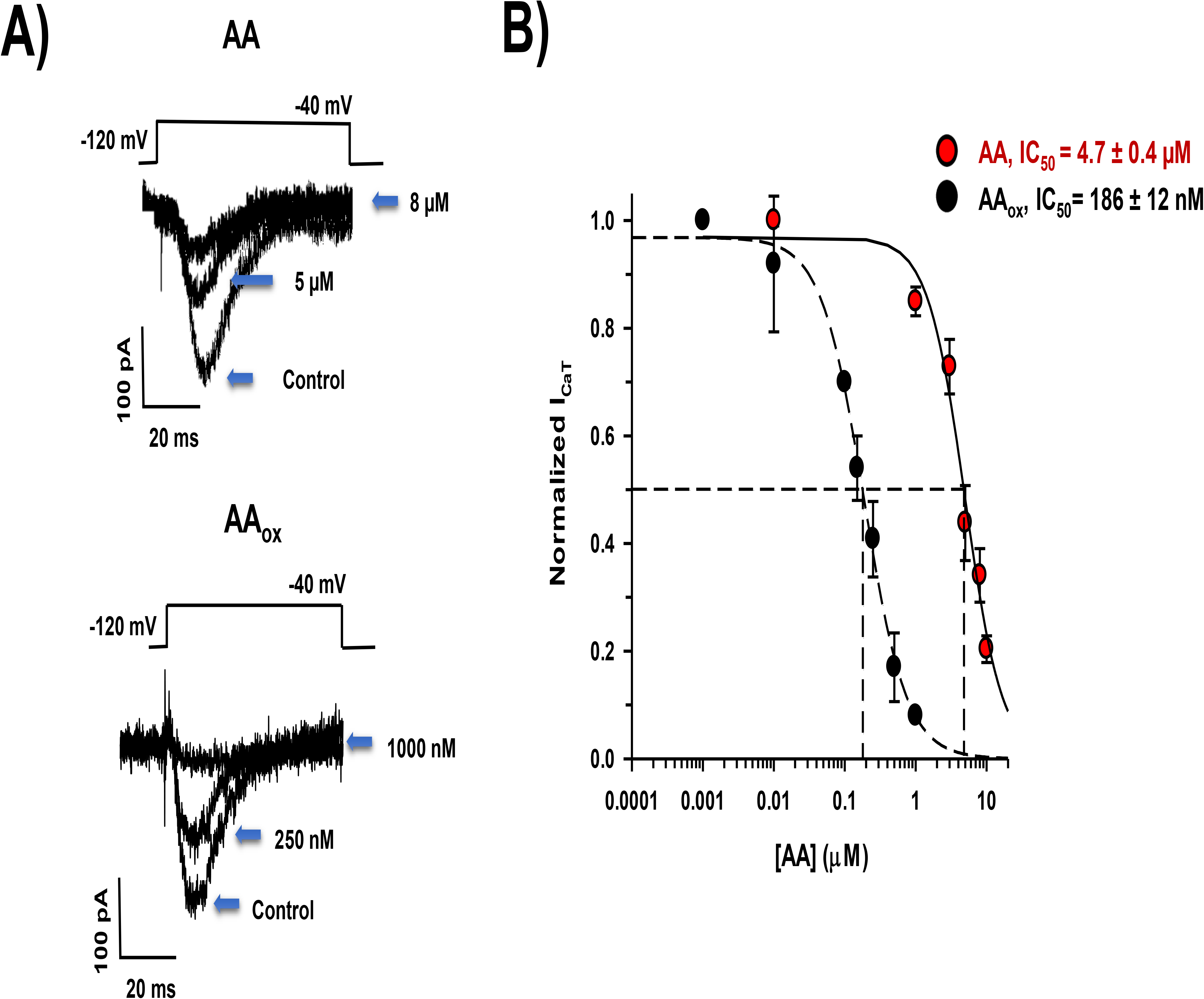
Oxidized arachidonic acid (AA_ox_) inhibits the spermatogenic cell T-type Ca^2+^ current (I_CaT_) more potently than non-oxidized AA. **A)** Representative traces evoked with a 200 ms depolarizing pulse at −40 mV from a −120 mV V_h_ in absence (control) or presence of different AA (upper traces) or AA_ox_ (lower traces) concentrations. **B)** Inhibition dose-response curves for AA (red circles) and AA_ox_ (closed circles). Following exposure to the indicated concentrations of AA or AA_ox_, I_CaT_ amplitude evoked with a −40 mV test pulse was reduced respect to the control at the same voltage. A smooth curve was generated with the Hill equation according to the parameters obtained from the I_CaT_ peak amplitude. The IC_50_ values were 4.7 ± 0.4 µM and 186 ± 12 nM; with Hill coefficients of 1.6 ± 0.3 and 1.4 ± 0.1 for AA and AA_ox_, respectively. In all cases, symbols represent the mean ± S.E.M. (n=4–6).

The spontaneous oxidation of plasma membrane phospholipids and FAs has been involved in mammalian infertility (Alvarez & Storey, 1995). Because there is a lack of information regarding the effects of AA_ox_, and they display a higher inhibitory potency (above shown) and differential effects on T-type Ca^2+^ channels, we decided to characterize the AA_ox_ effects on spermatogenic cell T-type Ca^2+^ currents.

### AA_ox_ does not affect neither time-to-peak nor inactivation kinetics of spermatogenic cell I_CaT_

The shift observed in the peak of the I-V curve could imply changes in I_CaT_ activation kinetics. In order to evaluate the AA_ox_ effect on I_CaT_ kinetics, we analyzed both the time-to-peak (t_p_) and the time constant of inactivation (τ_inac_) in the control condition and in the presence of AA_ox_. AA_ox_ products have no effect on inactivation or time-to-peak of spermatogenic cell T-type Ca^2+^ channels (Fig 4A and B). A previous report showed that fresh AA does not affect these kinetic parameters in heterologously expressed T-type Ca^2+^ channels (Talavera *et al,* 2004b). We corroborated that fresh AA (3 µM) did not change the time-to-peak or inactivation kinetics of spermatogenic cell I_CaT_ (Fig. EV3).

**Figure 4.**
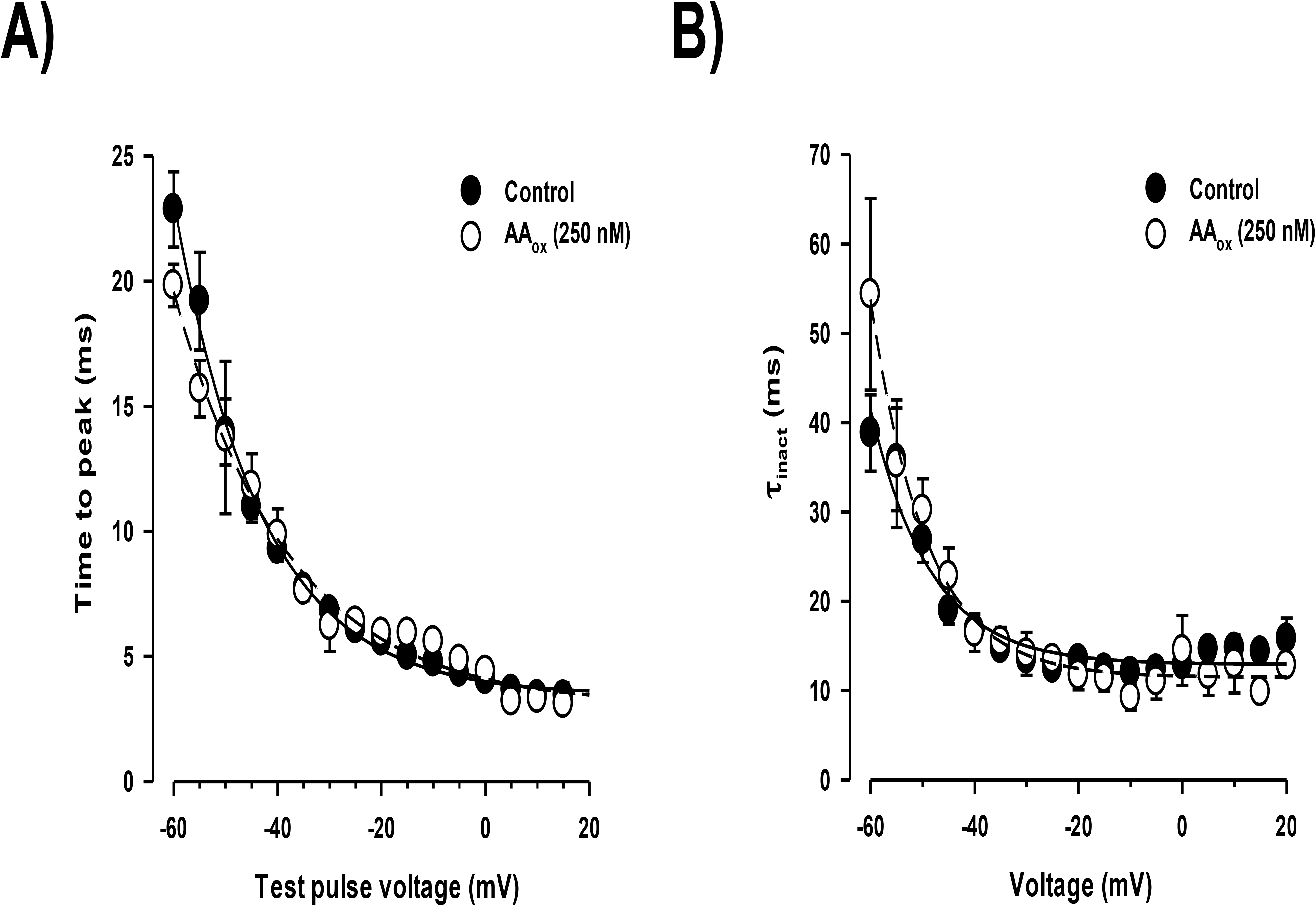
AA_ox_ neither influence time-to-peak nor the inactivation kinetics of spermatogenic cells I_CaT_. **A)** Voltage-dependence of I_CaT_ activation (time to peak). Addition of AA_ox_ (250 nM, open circles) does not alter the activation kinetics values compared to the control condition (closed circles) (n=4). Representative activation traces in absence (control, solid line) or presence of AA_ox_ (250 nM, dotted trace) are shown in lower panel. Time to peak values were obtained from current activation traces. **B)** I_CaT_ inactivation constant values (τ_inact_) were statistically similar after incubation with AA_ox_ (250 nM, open circles) compared to control conditions (closed circles). Representative inactivation traces in absence (control, solid line) or presence of AA_ox_ (250 nM, dotted trace) are shown in lower panel. τ_inact_ values were obtained from time constants of fitted current inactivation traces (n=5). In all cases, symbols represent mean ± S.E.M.

### AA_ox_ left shifts both the steady state inactivation and activation curves of I_CaT_

Since AA shifts the inactivation curve to more negative potentials, but does not affect the activation curve of heterologous express Ca_V_3.1 Ca^2+^ channels (Talavera *et al*, 2004), we tested how AA_ox_ affected steady-state inactivation (Fig. 5A) and activation (Fig. 5B) curves. In both cases, addition of AA_ox_ led to left shifts of the curves to more negative potentials, with changes of half inactivation voltage from V_50_= − 65 ± 1 mV in control to V_50_= − 74 ± 2 mV after AA_ox_ addition (Fig. 5A) and with changes of half activation voltage from V_50_= − 32 ± 0.5 mV in control to V_50_= − 43 ± 0.6 mV in the presence of AA_ox_ (Fig. 5B). As a consequence of both curve shifts, the I_CaT_ current window was also left shifted 10 mV and reached its maximum around −57 mV in presence of AA_ox_, and −47 mV in control conditions.

**Figure 5.**
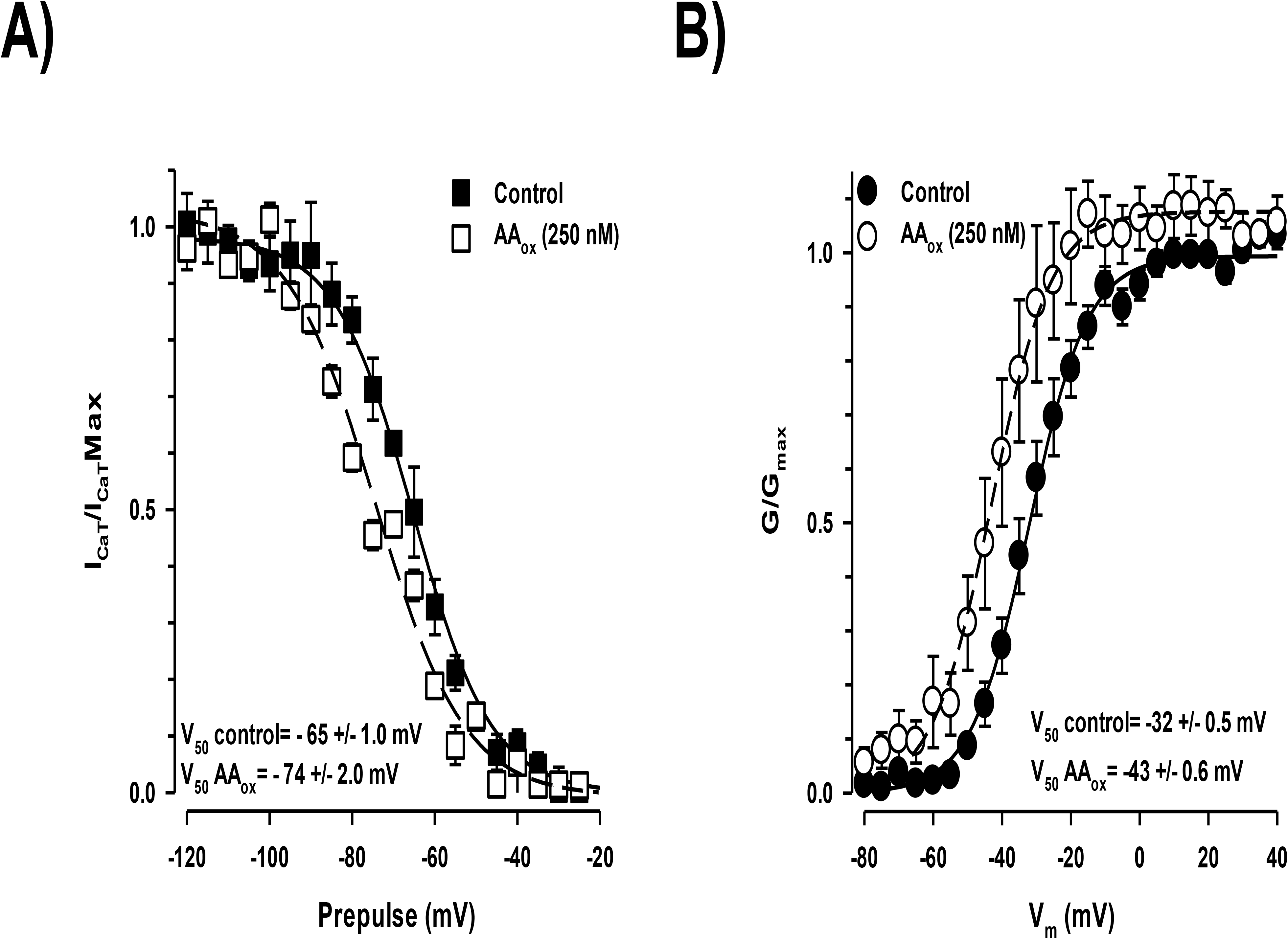
AA_ox_ left shifts both steady-state inactivation and activation curves of I_CaT_. **A)** Steady-state inactivation curves of I_CaT_ in control (closed squares) or experimental (AA_ox_, open squares, 250 nM) conditions. I_CaT_ were obtained in response to 1 sec preconditioning steps from −120 up to −20 mV followed by a depolarized test pulse to −40 mV. Peak current amplitudes obtained from a test pulse were normalized to the maximal current recorded from each cell and fitted to a Boltzmann relation. V_50_ inactivation values for control or in presence of AA_ox_ were − 65 ± 1.0 and −74 ± 2.0 mV, respectively (n=6). **B)** Voltage-dependent activation curves of I_CaT_ in control (closed circles) or experimental conditions (AA_ox_, open circles). I_CaT_ was evoked by a 30 ms depolarizing test pulses from a Vh of −120mV from −80 up to +40mV in 5 mV steps. Repolarization from test pulses was to −120 mV, and tail currents were normalized by the maximal current amplitude in each recording. V_50_ activation values for control or in presence of AA_ox_ were −32 ± 0.5 and −43 ± 0.6 mV, respectively (n=3–5). In all cases, symbols represent mean ± S.E.M. \**p*<0.05

### AA_ox_ slows spermatogenic cell I_CaT_ deactivation kinetics

A previous report indicates that fresh AA does not modify the deactivation of α_1G_ channels expressed in HEK cells (Talavera *et al*, 2004). To evaluate whether AA_ox_ regulates the I_CaT_ deactivation in spermatogenic cells, we applied a deactivation protocol in the absence or presence of AA_ox_. Deactivation of I_CaT_ was recorded and tail currents at different deactivating potentials from −110 to −60 mV, after a 5 ms activation pulse, were fitted by a single exponential function to calculated τ_Deact_. Deactivation time constant (τDeact) values of I_CaT_ were larger in the presence of AA_ox_ (Fig. 6 left panel, open circles) with respect to the control condition (Fig. 6 left panel, closed circles), indicating that AA_ox_ differentially affects spermatogenic cell I_CaT_.

**Figure 6.**
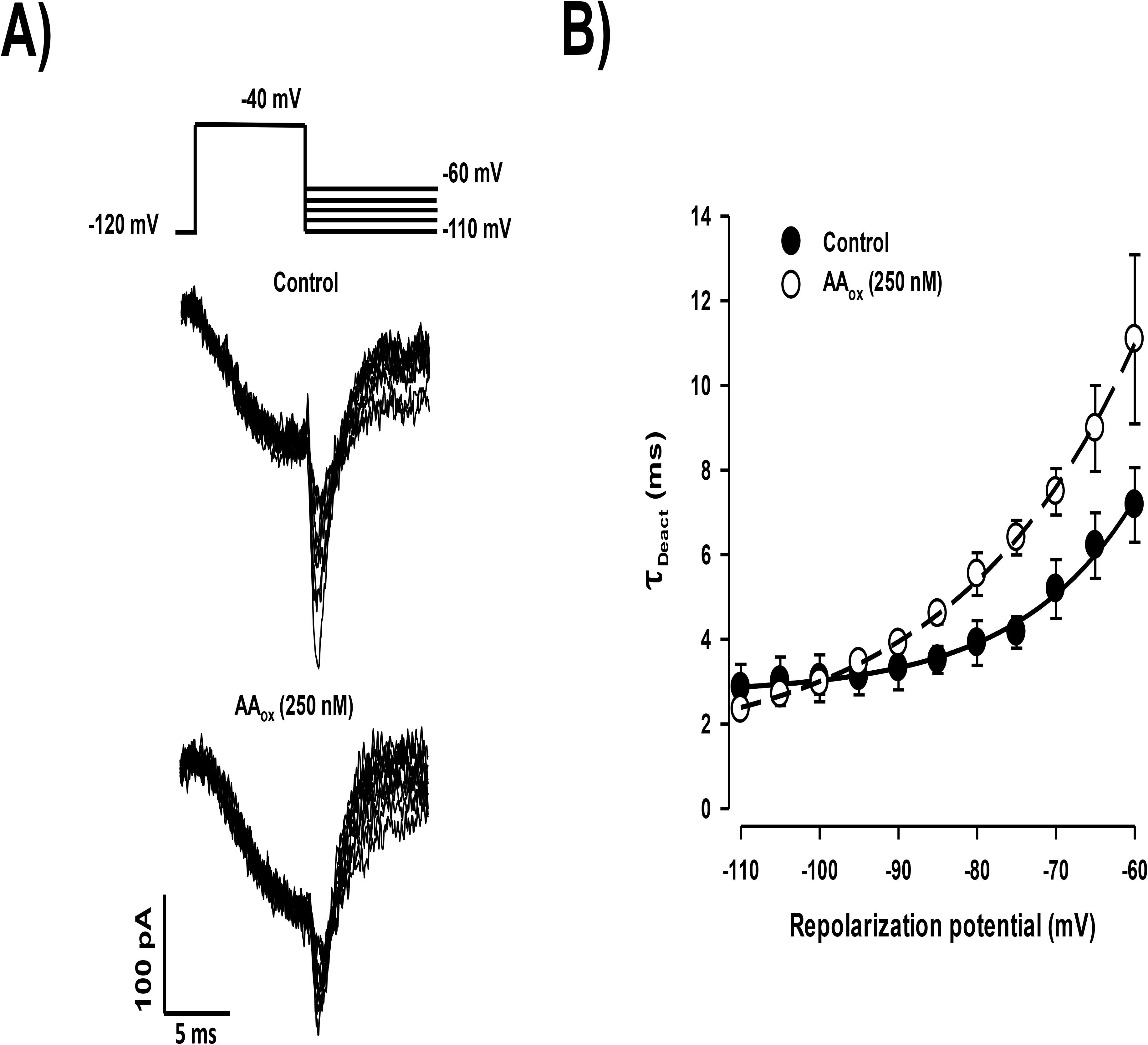
Deactivation kinetics of I_CaT_ is slower in the presence of AA_ox_. **A)** Representative tail current traces in absence (control, upper traces) or presence of AA_ox_ (lower traces). Tail currents at different deactivating potentials from −110 up to −60 mV after a 7 ms activation pulse were fitted by a single exponential function to calculate τ_Deact_ (n=4-6). For control, τ_Deact_ values were 2.9 up to 7.2 ms; for AA_ox_ treated spermatogenic cells, τ_Deact_ values were 2.1 up to 11.4 ms. The voltage protocol is shown above current traces, V_h_ was −120 mV. **B)** Voltage-dependence of I_CaT_ deactivation constant (τ_Deact_) in control (closed circles) or presence of AA_ox_ (open circles). In all cases, symbols represent mean ± S.E.M. (n=4-6) \**p*<0.05

### I_CaT_ recovery from inactivation is delayed by AA_ox_

Taking into account that AA does not affect the recovery from inactivation of α_1G_ Ca^2+^ currents when applying a V_h_ more negative than −100 mV (Talavera *et al*, 2004), we evaluated the effect of AA_ox_ on this parameter of spermatogenic cell I_CaT_. Representative traces of recovery from inactivation show that AA_ox_ significantly increases the required time for channel recovery from inactivation state from τ_Rec(control)_ = 125 ±7.0 ms in control (Fig. 7A and B, closed circles) to τ_Rec(AAox)_ = 172 ±3.0 ms (Fig. 7B, open circles).

**Figure 7.**
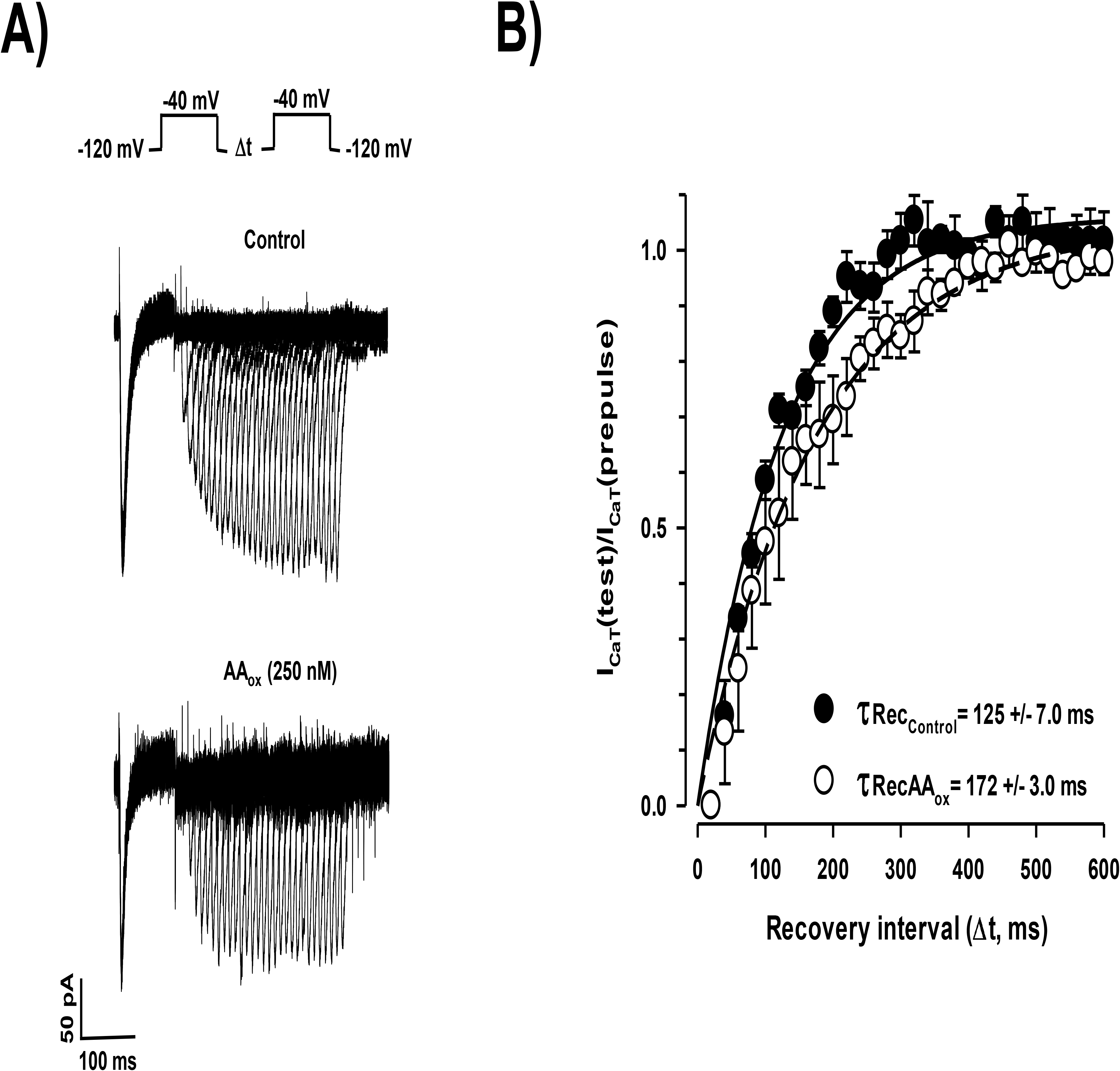
Recovery from inactivation of I_CaT_ is slowed by AA_ox_. **A)** Representative current traces for control (upper traces) and experimental (AA_ox_ 250 nM, lower traces) conditions. I_CaT_ was obtained using a two-pulse protocol, where the peak current amplitude evoked by the test pulse at −40 mV was divided by the Ca^2+^ current amplitude evoked by the prepulse separated by different lapse times, V_h_ was −120 mV. **B)** Recovery from inactivation curves of I_CaT_ in control (closed circles) or presence of AA_ox_ (open circles). Normalized I_CaT_ was obtained using a two-pulse protocol described above. Temporal course of recovery from inactivation was fitted by a single exponential function and the recovery from inactivation half time (T_50_ rec) was calculated in absence (control) or presence of AA_ox_ with values of 125 ± 7 and 172 ± 3 ms, respectively (n=4–6). In all cases, symbols represent mean ± S.E.M. \**p*<0.05

### Albumin reverts AA_ox_-induced I_CaT_ inhibition, but reversion was both AA_ox_ concentration and time incubation dependent

It is know that AA-induced inhibition of α_1G_ Ca^2+^ currents is rapidly reversed upon washing with BSA (Talavera *et al*, 2004). As previously reported, external BSA increases the I_CaT_ amplitude of spermatogenic cells (Espinosa *et al*, 2000; López-González *et al*, 2016). Consistently, we observed that perfusing spermatogenic cells with a BSA (1%) solution increased the I_CaT_ amplitude around 60% with respect to the control in the first 2 minutes of perfusion. Thereafter, the current amplitude decreased, and a plateau level was established being 40% higher than the control condition (Fig. 8A, gray vs closed circles). Considering that BSA is an excellent AA carrier (Brash, 2001) and also acts as a very potent protector against oxidized lipids in sperm due to its high affinity for them (Alvarez & Storey, 1995), we evaluated if this protein could protect the I_CaT_ in spermatogenic cells from oxidized AA. As a first approach to explore this possibility, we used AA_ox_ preincubated with BSA (1%), in an equimolar stoichiometry. Adding an AA_ox_/BSA (1%) containing solution eliminated the I_CaT_ amplitude increase and produced a partial inhibition (∼50%) probably due to the transference of AA_ox_ into the spermatogenic cell plasma membrane (Fig. 8B, dark gray circles). This inhibition percentage was statistically similar to the observed inhibition in presence of a lower AA_ox_ concetration (250 nM; Fig. 8B, open circles). Interestingly, the AA_ox_/BSA-induced I_CaT_ inhibition was fast and spontaneously reverted in spermatogenic cells (Fig. 8B, dark gray circles).

**Figure 8.**
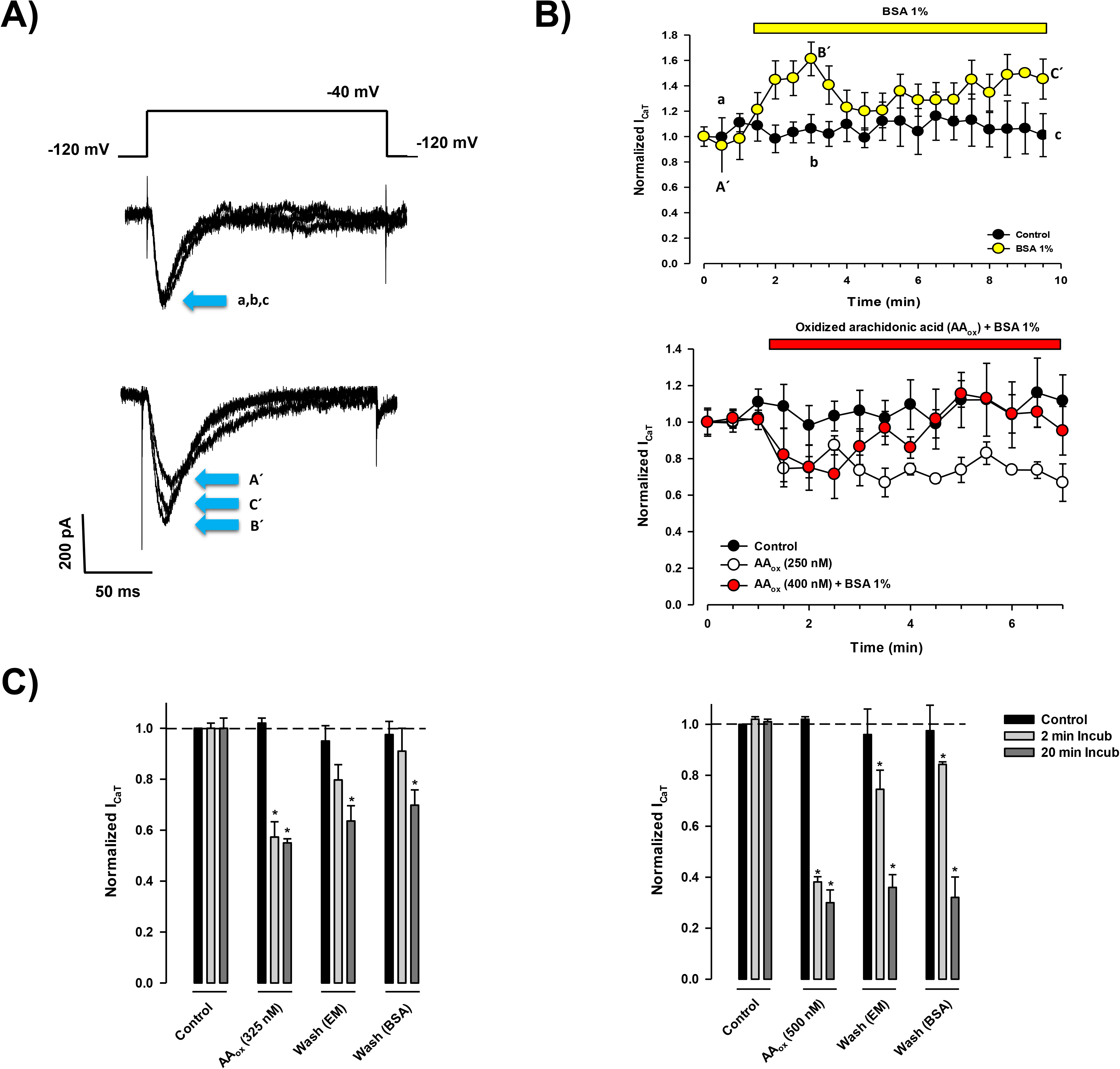
Albumin increases the I_CaT_ amplitude and reverts the AA_ox_-induced inhibition of spermatogenic cell Ca^2+^ current. **A)** Representative traces evoked with a 200 ms depolarizing pulse at −40 mV from a V_h_ −120 mV in absence (control, upper traces) or presence BSA (1%, lower traces) at 0.5 (A’, a), 3 (B’,b) or 10 (C’,c) min after the first evoked trace (0 min). **B)** Upper panel: normalized I_CaT_ temporal course from electrophysiological recordings showed in panel A. In control condition, I_CaT_ rundown was less than 5% (closed circles). Perfusing a BSA solution (1%, open bar) rapidly increased the I_CaT_ amplitude (open circles). Lower panel: addition of AA_ox_ (250 nM, open circles) inhibited around 40% the normalized I_CaT_ within 0.5-min lapse, compared to control (closed circles). Maximal AA_ox_ inhibition was stable through time (10 min). However, preincubation of a higher AA_ox_ concentration (400 nM) with BSA (1%), in an equimolar stoichiometry, allowed complete reversion of I_CaT_ inhibition (red circles). **C)** The AA_ox_-induced inhibition of I_CaT_ was concentration and time dependent. Left panel: Addition of AA_ox_ (325 nM) inhibited the normalized I_CaT_ amplitude around 43%. This inhibition percentage was statistically similar after 2 or 20 min of incubation. Washing AA_ox_ with external solution (EM) recovered the I_CaT_ current amplitude up to 79% after a short incubation (2 min); whereas after a long-lasting AA_ox_ inhibition (20 min), I_CaT_ was recovered up to 63% of its maximal amplitude at the same concentration. BSA (1%) solution improved the AA_ox_ wash out from spermatogenic cells; where we observed an I_CaT_ amplitude recovery up to 90% after 2 min-lasting AA_ox_ incubation vs 70% of I_CaT_ amplitude recovery after 20 min of AA_ox_ incubation. Right panel: A higher AA_ox_ concentration (500 nM) inhibited a bigger fraction of the I_CaT_ amplitude (62%), and this inhibition was statistically similar at 2 or 20 min of incubation. Perfusing spermatogenic cells with external solution (EM) reverted more efficiently the I_CaT_ inhibition induced by an AA_ox_ short incubation (2 min, 75% of I_CaT_ amplitude recovery) than a long-lasting AA_ox_ inhibition (20 min, 63% of I_CaT_ inhibition). BSA (1%) solution slightly improved the AA_ox_ wash out; where we observed an I_CaT_ recovery up to 82% after 2 min-lasting AA_ox_ incubation but no I_CaT_ recovery after long-lasting AA_ox_ incubation (20 min). In all cases, symbols or bars represent the mean ± S.E.M. (n=4). \**p*<0.05.

The reversibility of AA_ox_ inhibitory effect on T-type Ca^2+^ currents of spermatogenic cells was studied under various conditions. Cells were incubated in the absence (control) or presence of 325 (Fig. 8C, left panel) or 500 nM (Fig. 8C, right panel) of AA_ox_, for 5 or 20 min-lasting incubations for both concentrations. After incubation with AA_ox_, cells were subsequently perfused with external media or with BSA-containing media. Our results showed that the reversibility of AA_ox_ inhibitory effect on T-type current depends on both incubation time and concentration. For instance, the recovery of T-type Ca^2+^ current amplitude (∼80%) after 2 min of AA_ox_-induced inhibition in presence of 325 or 500 nM showed no statistical difference when it was washed with either BSA-free or BSA-containing extracellular media (Fig. 8C, gray bars). On the contrary, the reversibility of the T-type Ca^2+^ current inhibition after 20 minutes of incubation with 325 nM AA_ox_ was partial (60%) but it was irreversible in presence of 500 nM AA_ox_, even when exposed to BSA-containing media (Fig. 8C, dark gray bars).

## DISCUSSION

Various groups have reported deleterious effects of spontaneous oxidation of plasma membrane phospholipids and FAs in mammalian sperm. In general, phospholipids and FAs oxidation of mammalian sperm produces plasma membrane damage (Jones & Mann, 1973, 1976, 1977, Jones *et al*, 1978, 1979; Alvarez & Storey, 1995), a reduction in antioxidant activity, spermatic abnormality, apoptosis, and DNA fragmentation (Chabra *et al*, 2014; Mohamed & Mohamed, 2015). This multifactorial phenotype is a consequence of different oxidized plasma membrane compounds which regulate a diversity of intracellular molecular targets.

Here we focused on the specific effect of non-enzymatically oxidized AA products on T-type Ca^2+^ channels of spermatogenic cells, considering the physiological relevance of both I_CaT_ and AA in the male germ cell line (Sánchez-Cárdenas *et al*, 2012; Paillamanque *et al*, 2016, 2017). Furthermore, there are few reports that show the functional modification of spermatogenic cells, by secreted molecules from Sertoli cells, especially at the meiotic and post-meiotic stages (reviewed in Paillamanque *et al*, 2016). Even less is known about the early effects of specific oxidized FAs on ion channels in spermatogenesis.

As a first approach to characterize the early effects of oxidized FAs on spermatogenic cells physiology, we induced the non-enzymatic oxidization of AA and obtained two main oxidized products which seem to be in equilibrium (Fig. 1). Unlike FAs which are oxidized by enzymes with a high degree of positional and conformational specificity, unsaturated FAs are also vulnerable to many types of oxidation by non-enzymatic mechanisms that result in the formation of many different oxidized forms. Our findings indicate that non-enzymatic oxidation of AA continuously produced AA_ox_ products, two of which seem to be in constant equilibrium, precluding their independent evaluation. Therefore, the inhibition of spermatogenic cell I_CaT_, here reported, must be caused by the mixture of both AA_ox_ products. According to MS and NMR spectra, the AA_ox_ products lose their four double bonds and all of them were completely hydroxylated, whereas they could contain at least one epoxide group at position 8 or 11 (Fig. 1).

### AA and AA_ox_ inhibit *I_CaT_* by different mechanisms

In the present study we investigated the mechanism of inhibition of spermatogenic cell T-type current by AA_ox_ as an essential step in understanding the influence of oxidized FAs on the physiology of male germ cell line. We show that AA_ox_ inhibits I_CaT_ more potently than non-oxidized AA (Fig. 2). The majority of earlier reports indicate that AA regulates both α_1G_-and α_1H_-encoded Ca^2+^ currents in the micromolar range (Zhang *et al*, 2000; Chemin *et al*, 2002; Talavera *et al*, 2004). Regardless of the Ca^2+^ current type (N, L, or T) and cell type (rev in Meves, 2008), the estimated IC_50_ was between 3 and 10 µM. Our results confirmed that AA inhibits spermatogenic cell I_CaT_ in the same concentration range (IC_50_ = 4.7 µM); however, AA_ox_ showed a significantly higher inhibitory potency (IC_50_ = 186 nM).

Even though the inhibition of T-type Ca^2+^ channels by AA has been described in several studies, the contribution of its metabolites is not fully understood. Some reports suggest that Ca^2+^ current inhibition is caused by AA itself. For instance, Zhang *et al* (2000) showed that application of lipoxygenase (nordihydroguaiaretic acid) and cyclooxygenase (indomethacin) inhibitors does not prevent the AA-induced inhibition of α_1H_-encoded Ca^2+^ current, indicating a direct inhibition by AA. Results of Talavera *et al* (2004) and Schmitt & Meves (1995) support this hypothesis since addition of 17-ODYA, an AA-derivate metabolite, shows no effect on α_1G_ currents. Furthermore, these authors state that ETYA, a non-metabolizable analogue and inhibitor of the AA metabolism, did not influence the inhibition by AA, suggesting that AA and not its metabolites, is directly involved in T-type Ca^2+^ current regulation. However, other reports suggest the AA metabolites do participate in Ca^2+^ current inhibition. Addition of 8,9-EET, a specific cytochrome P-450 produced metabolite, together with AA led to 31% reduction of G_max_, suggesting that the epoxygenase metabolite of AA (8,9-EET) can be partially involved in the inhibition of α_1H_-encoded channel activity. Interestingly, ETYA by itself led to partial inhibition of α_1G_ current even though its potency was significantly lower than that of AA (Talavera *et al*, 2004). Contrary to the above mentioned studies, our findings indicate non-enzymatically produced AA_ox_ compounds inhibit spermatogenic T-type Ca^2+^ by acting themselves. They can be easily removed from spermatogenic cell plasma membranes by washing which is mildly enhanced by including BSA (Fig. 8).

The effects of AA_ox_ on some biophysical parameters of I_CaT_ were clearly different from those previously reported for AA. For instances, non-oxidized AA induces a decrease of the current amplitude but does not significantly modify the I-V curve shape (Talavera *et al*, 2004); whereas AA_ox_ shifted the peak of I-V curve around 10 mV to more negative potentials (Fig. 3). AA shifts the steady-state inactivation curve to more negative potentials, but it does not affect the activation curve (Talavera *et al*, 2004). AA-induced Ca^2+^ current inhibition has been explained as a consequence of a decrease in channel open probability (Liu & Rittenhouse, 2000), which causes an increase in the number of channels in a non-conducting conformation, that is in either closed or inactivated states (Meves, 2008). In contrast, AA_ox_ left shifted both the activation and steady-state inactivation curves (Fig. 5). Whereas AA does not modify the deactivation of T-type Ca^2+^ channels (Talavera *et al*, 2004), I_CaT_ deactivation kinetics was slower in presence of AA_ox_ (Fig. 6). Lastly, AA does not affect recovery from inactivation with holding potentials more negative than −100 mV (Talavera *et al*, 2004) while AA_ox_ significantly increased the time for recovery from inactivation of I_CaT_ at a holding potential of −120 mV (Fig. 7).

Altogether, these data indicate that the molecular mechanism of the AA_ox_-induced inhibition of I_CaT_ is different from that proposed for AA. In the case of AA_ox_, the activation threshold of I_CaT_ I-V curve and the activation curve were both left shifted; this effect could be explained by an increase in the transitions between the closed states of the activation pathway. Our results indicate AA_ox_ slowed the deactivation kinetics of I_CaT_, suggesting that it does affect the transition from the open to the nearest closed state. Thus, in presence of AA_ox_, spermatogenic cell T-type Ca^2+^ channels could transit from the open to inactivated states and spend more time inactivated rather than going back to close states. Indeed, AA_ox_ profoundly affected the inactivation properties of spermatogenic cell I_CaT_. AA_ox_ shifted the voltage dependence of steady-state inactivation by 10 mV to more negative potentials and slowed the kinetics of the recovery from inactivation. Therefore, one of the most important conclusions in this report is that in presence of AA_ox_, the fraction of inactivated Ca^2+^ channels of spermatogenic cells is increased at voltages where they are usually not inactivated and their recovery from inactivation is slowed. The change in both processes related to Ca^2+^ channel inactivation reduce both the fraction of Ca^2+^ channels able to open and their contribution to the macroscopic current amplitude, therefore reducing the Ca^2+^ current.

### AA_ox_-induced I_CaT_ inhibition may occur in a membrane-delimited manner

As previously mentioned, it has been argued that AA inhibits α_1G_ channels partitioning into the plasma membrane (Talavera *et al*, 2004). When added to cell-free inside-out patches, Ca^2+^ current inhibition by AA was associated to the compound itself, as its metabolites could not be produced by enzymes given the lack of intermediary reagents in the bath solutions (Arreaza *et al*, 1997). Furthermore, AA-induced inhibition of Ca^2+^ current was removed within 0.5 min with a BSA-containing external solution suggesting AA was inhibiting Ca^2+^ channels in a membrane-delimitated manner (Talavera *et al*, 2004), though the comparison washing with only media was not performed.

Our results showed that preincubation of AA_ox_ with BSA, in an equimolar manner, reduced the AA_ox_-induced I_CaT_ inhibition which then reverted (Fig. 8). As previously mentioned, albumin is a very potent protector against lipid peroxidation in sperm due to its high affinity for oxidized FAs, and also acts as an excellent AA carrier (Alvarez & Storey, 1995; Brash, 2001). Our preincubation assays could suggest that AA_ox_ is incorporated to the plasma membrane of spermatogenic cells as a first step, then the oxidized FA reaches a partition equilibrium between the plasma membrane and extracellular free-BSA, recovering the I_CaT_ amplitude. To better understand these effects, we evaluated the effect of washing with media ±BSA at different AA_ox_ concentrations and incubation times. Our findings showed that the reversibility of AA_ox_ inhibitory effect on I_CaT_ depends on both AA_ox_ incubation time and concentration. For instance, Ca^2+^ current amplitude recovery after 2 minutes of 325 or 500 nM AA_ox_-induced inhibition was statistically similar washing with extracellular media ± BSA. In contrast, I_CaT_ reversibility after 20 minutes of incubation with 325 nM AA_ox_ was partial, but it was completely irreversible at 500 nM AA_ox_, even after washing with a BSA-containing media. It is likely that when incubating 20 min with 500 nM AA_ox_, a significant percentage of the oxidized FAs flip to the internal lipid layer precluding their removal from plasma membrane by the BSA containing solution and preventing Ca^2+^ current amplitude recovery during washing. This result is consistent with the proposal that at high AAox concentration and incubation times inhibition of I_CaT_ is at least partially membrane delimitated. Indeed, washing spermatogenic cells with BSA-containing media slightly improves the recovering of I_CaT_ amplitude from a AA_ox_ short-time inhibition.

As previously reported, external BSA increases the I_CaT_ amplitude of spermatogenic cells (Espinosa *et al*, 2000; López-González *et al*, 2016). Consistently, we observed that perfusing spermatogenic cells with a BSA solution increased the I_CaT_ amplitude around 40% higher than the control condition (Fig. 8). This result suggests BSA could increase the I_CaT_ amplitude by removing the tonic inhibition from a fraction of Ca^2+^ channels by endogenous AA/AA_ox_ present in the spermatogenic cell plasma membrane.

## MATERIALS AND METHODS

### Fresh (AA) and oxidized arachidonic acid (AA_ox_) preparation

The sodium salt of Arachidonic acid from Mortierella alpina (AA) was ordered from SIGMA-ALDRICH (SML1395) and dissolved according to instruction with 100% Ethanol to a final concentration of 150 mM (50 mg/ml). Part of freshly prepared AA was used for analyzes and another part was saved for preparation of non-enzymatically oxidized sample. For oxidation, 5 mg of AA dissolved in 100 µL of ethanol were placed in a glass flask with light protection and stirred for 24 h at room temperature (∼24°C) aerating the flask for 30 s each 4-5 hours. Thereafter, the volume of the sample was checked and adjusted to the initial level (100 µL) to preserve the calculated concentration (150 mM). The samples obtained were characterized using HPLC and peaks were separated to isolate AA_ox_ forms.

### Purification of AA_ox_ products by reverse phase HPLC

For HPLC, 10 µL of 150 mM AA_ox_ was dissolved in 400 µL 25% Acetonitrile + 0.1% TFA solution and vortexed for 30 s until sample become transparent. Shortly after vortexing, peaks that corresponds to oxidized AA forms were separated by rpHPLC using an analytical CN column (Varian Microsorb-MV 100 CN, 250 x 4.6 mm) thru a gradient from solvent A (0.1 % TFA in water) to solvent B (0.1 % TFA in acetonitrile) starting with a 30:70 ratio of solvent B to A, respectively with then further increasing the B solvent concentration 1%/min. The flow rate was 1mL/min. Effluent absorbance was monitored at 230 nm Components corresponding to fractions p42 and p45 were dried under vacuum to prevent further decomposition and dissolved in ethanol. These fractions were tested for their effect on T-type Ca^2+^ currents in spermatogenic cells and re-purification. For each peak, procedure of separation, drying and testing was performed at least 3 times.

### Mass spectrometry of AA_ox_ products

The HPLC-MS experiments were performed on a HPLC Agilent 1260 Infinity coupled to an MS-Q-TOF 6530 with Jet Stream ESI using a C18 column PoroShell 120EC-C18 2.1X100 mm 2,7 µm (Flores-Solis *et al*, 2016). For each analysis, 10 µL of oily sample were dissolved in 20 mL of 60/40 methanol/water solution. Isocratic conditions were used as solution to dissolve the sample. HPLC chromatograph and mass spectra are shown in Fig. EV1.

### NMR experiments

NMR samples were prepared with CDCl3 (99.99 % D). ^1^H NMR experiments were acquired in a 500 MHz Varian Inova and 300 MHz Bruker Advance at 297 K. NMR spectra of arachidonic acid was used directly from Aldrich. TMS was used as internal standard.

### Electrophysiology

Spermatogenic cells were collected from CD1 mouse and prepared as described previously (López-González *et al*, 2016). After disaggregation procedure spermatogenic cells were placed on the stage of an inverted microscope (Diaphot 300, Nikon) and kept there for 5 min to adhere to the coverslip surface. Membrane currents were recorded with an Axopatch 200B amplifier (Molecular Devices, Sunnyvale, CA). Macroscopic Ca^2+^ currents were acquired at a sampling frequency of 20 kHz (50 ms) and filtered at 5 kHz (four-pole Bessel filter) during recording, and then digitized using a Digidata 1440A interface (Molecular Devices). Linear capacitive currents were minimized analogically using the capacitative transient cancellation feature of the amplifier. All experiments were carried out at room temperature (22°C) and the holding potential (HP) was −120 mV. Borosilicate glass was used for preparation of patch pipettes by pulling with a laser micropipette puller P-2000 (Sutter Instruments Co., Novato, CA). The typical micropipette electrical resistance was 6–8 MΩ when filled with internal solutions. The recording extracellular (EM) solution contained (in mM): 125 TEACl, 5 CaCl_2_, 10 N-2-hydroxyethylpiperazine-W-2-ethanesulfonic acid (HEPES) and 10 D-glucose, pH 7.3 adjusted with TEAOH. Intracellular solution contained (in mM): 120 CsMeSO_4_, 10 EGTA, 5 MgCl_2_, 10 HEPES, 10 D-glucose and 60 glutamic acid, pH 7.4 was adjusted with CsOH. Osmolarity of external (290 mOsmol/kg) and internal (265 mOsmol/kg) media was monitored using a vapor pressure osmometer (Wescor). Obtained data were analyzed with the Clampfit 10.7 (Molecular Devices, Sunnyvale, CA).

### T-type Ca^2+^ currents analysis

Current to voltage relationships (I–V curves) were obtained by plotting the peak amplitude of Ca^2+^ currents as a function of their respective membrane potential during the test pulse. Dose-response curves were fitted using the Hill equation:

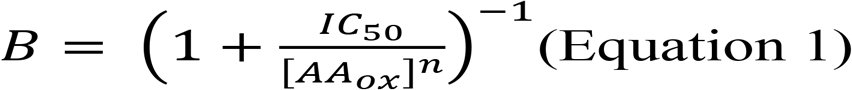

where *B* is the normalized blocked current, *IC*_*50*_ is the AA_ox_ concentration giving half-maximal inhibition, and *n* is the Hill coefficient. Steady-state inactivation and activation curves were obtained from the ratios of peak current amplitudes normalized to the maximal current amplitude by fitting to the following Boltzmann relation:

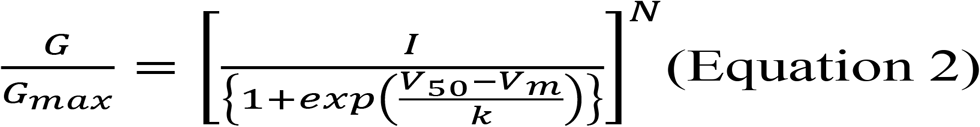

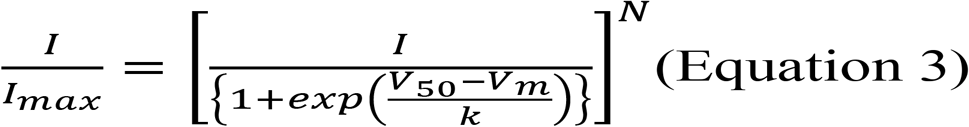

where *G*_*max*_ is the maximal conductance, *I*_*max*_ is the maximal Ca^2+^ current amplitude, *V*_*50*_ is the voltage at which half of the current is activated or inactivated, *V*_*m*_ is the membrane potential, *k* is the slope factor, and *N* is the power factor. Time constants for activation and inactivation were measured by fitting individual traces to the following kinetic equation:

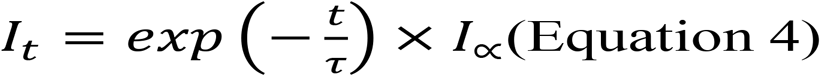

where *t* is time, τ is the activation (τ_act_) or inactivation (τ_inact_) time constant, and *Iα* is an amplitude scaling factor. For the deactivation analysis, data were obtained by fitting a curve to the tail-currents following settling of 95% of the membrane capacitance transient. In all cases, tail-currents were well fitted by a single exponential equation (equation 4) to determine their fast component.

### Statistical data analysis

Statistical analysis was performed using the Sigmaplot 12.3 software (Systat Software, Inc). Data were expressed as the mean ± standard error of the mean (S.E.M.). Statistical significance was determined using Student paired t-test or Analysis of variance (ANOVA) and Tukey’s test for multiple comparisons. A *p*<0.05 was considered significant. All the experiments were repeated at least three times.

## ACKNOWLEDGEMENTS

The authors thank Shirley Ainsworth, for technical assistance; Juan Manuel Hurtado, Roberto Rodríguez, Omar Arriaga and Arturo Ocádiz for computer services; and Dr. Otto Geiger for his expertise and assistance in chromatography. OB was depositary of a fellowship from Dirección General de Asuntos del Personal Académico/ Universidad Nacional Autónoma de México (DGAPA/ UNAM). This work was supported by CONACyT-México (71 and 128566 to AD and 84362 to ILG) and DGAPA/ UNAM (IN205516 to AD and IN205518 to ILG). In its final stage, this work was supported by the Ministry of Education, Youth and Sports of the Czech Republic - projects “CENAKVA” (No. CZ.1.05/2.1.00/01.0024), “CENAKVA II” (No. LO1205 under the NPU I program), CZ.02.1.01./0.0/0.0/16_025/0007370 Reproductive and genetic procedures for preserving fish biodiversity and aquaculture and by the Grant Agency of the University of South Bohemia in Ceske Budejovice (125/2016/Z) and by the Czech Science Foundation (No. 18-12465Y).

## AUTHORS’ CONTRIBUTIONS

ILG and AD conceived and coordinated the study, and contributed to the analyses of the experiments and to write the final version of the manuscript. OB and ILG acquired and analyzed electrophysiological data. HPLC separation of oxidized compounds was conducted by OB, FLS and GC. FDP acquired and analyzed the MS and NMR data. All authors reviewed the results and approved the final version of the manuscript.

## EXPANDED VIEW FIGURE LEGENDS

**Figure EV1. Mass spectra of three AA_ox_ products separated by HPLC**

**A)** Methylation was made before GC-MS analysis. Fraction 38 (p38, upper panel) differs from fractions 42 (p42, middle panel) and 44 s (p44, lower panel). **B)** Peak fragments of fractions 42 and 44 are the same (p42 vs p44, middle and lower panels) and differ from fragments of fraction 38 (p38, upper panel). These results confirm that p42 and p44 fractions are very similar compounds.

**Figure EV2. Arachidonic acid reduces the I_CaT_ amplitude but does not displace the I-V curve peak current**

**A)** Representative current-voltage relationships obtained with 200 ms test pulses of 10 mV steps in the −80 to +40 mV range from a V_h_ of −120 mV. Incubation with AA (5 µM, open circles) inhibited the I_CaT_ amplitude current around 60% with respect to the control (closed circles). **B)** Voltage of the I-V curve peak in absence (control: −40 ± 2 mV, closed bar) or presence of AA (5 mM: −40 ± 1 mV), respectively. In all cases, symbols represent the mean ± S.E.M. (n=4).

**Figure EV3. AA does not alter time-to-peak or the inactivation kinetics of spermatogenic cells I_CaT_**

**A)** Voltage-dependence of I_CaT_ activation (time to peak). Addition of AA (3 µM, open circles) does not alter the activation kinetics values compared to the control condition (closed circles) (n=4). **B)** I_CaT_ inactivation constant values (τ_inact_) were statistically similar after incubation with AA (3 µM, open circles) compared to control conditions (closed circles). In all cases, symbols represent mean ± S.E.M. (n=4).

